# Differential roles for IL-4Rα and IL-13Rα1 in immune cell infiltration and epithelial remodeling in experimental eosinophilic gastritis

**DOI:** 10.64898/2026.01.15.699676

**Authors:** Anish Dsilva, Shraddha Sharma, Shireen Barakey, Michal Itan, Ariel Munitz

## Abstract

**Rationale:** Eosinophilic gastritis (EoG) is a chronic inflammatory disease characterized by infiltration of eosinophils and mast cells, epithelial remodeling, and fibrosis. Although EoG is increasingly recognized as a distinct type 2 inflammatory disease, the cellular and molecular events that drive disease pathogenesis remain poorly understood. This is due in part to the absence of robust and physiologically relevant experimental models that recapitulate human disease.

**Methods:** Experimental EoG was induced in wild type and *Il13ra1^-/-^*mice by repeated intragastric oxazolone challenges in skin-sensitized mice. IL-4Rα1 was neutralized using antibodies. Gastric histopathology was determined by H&E, anti-Ki67, chloroacetate esterase and anti-MBP staining. Gastric RNA was subjected to RNA sequencing.

**Results:** Experimental EoG resulted in robust gastric eosinophilia, mastocytosis, epithelial remodeling, and subepithelial fibrosis. Transcriptomic profiling of gastric tissue revealed broad upregulation of type 2 cytokine and epithelial-remodeling genes, including *Il4ra*, *Il4i1, Ccl5, Muc4, Mmp10*, *Mcpt1/2, Areg, Pparg* and *Tff2*. The transcriptome profile of experimental EoG was markedly distinct from that of experimental EoE despite both models being initiated by oxazolone, suggesting that identical inflammatory triggers elicit tissue-specific and context-dependent transcriptional programs. Blockade of IL-4Rα signaling abrogated both eosinophil and mast cell infiltration and attenuated epithelial remodeling, whereas genetic deletion of *Il13ra1* selectively suppressed epithelial remodeling without affecting inflammatory cell recruitment.

**Conclusion:** These findings establish experimental EoG as a robust model for dissecting the cellular and molecular mechanisms driving gastric type 2 inflammation. They further define receptor-specific roles for IL-4Rα and IL-13Rα1 in coordinating immune infiltration and epithelial remodeling in EoG.

## Introduction

Eosinophilic gastrointestinal disorders (EGIDs) are a group of increasingly recognized conditions characterized by tissue-specific eosinophilic inflammation of the gastrointestinal tract^1,2^. Eosinophilic gastritis (EoG) is a chronic inflammatory disease that belongs to the broader spectrum of non-esophageal EGIDs. It is histologically defined by dense infiltration of eosinophils and mast cells within the gastric mucosa, accompanied by epithelial hyperplasia, subepithelial fibrosis, and architectural remodeling^3^. Clinical symptoms of EoG include nausea, abdominal pain, early satiety, and dyspepsia^3^. Despite growing prevalence and clinical recognition, the pathophysiological mechanisms underlying EoG remain poorly defined.

Accumulating clinical and translational evidence supports an allergic disease etiology for EoG^4^. A substantial proportion of patients exhibit comorbid atopic conditions, including allergic rhinitis, asthma, food allergy^5^. Like other allergic diseases and EGIDs, including eosinophilic esophagitis (EoE), EoG is associated with elevated expression of type 2 cytokines^6^. These include interleukin (IL)-4, IL-5, and IL-13, as well as eosinophil-attracting chemokines including CCL26 (eotaxin-3), which correlate with the degree of tissue eosinophilia^6^. Furthermore, case studies and retrospective analyses suggest that treatment of EoG with anti-IL-4Rα (i.e., dupilumab) may be beneficial^7–9^. These observations implicate type 2 cytokine signaling in EoG pathogenesis. However, the functional contribution of individual cytokines and/or their downstream signaling pathways in EoG has not been systematically examined.

Insights into type 2 cytokine-driven tissue pathology in EGIDs, have largely emerged from studies of EoE, the most extensively characterized EGID^10^. In EoE, IL-13 has been identified as a critical effector cytokine that drives epithelial remodeling, basal cell hyperplasia, barrier dysfunction, and fibrosis, positioning the epithelium as a central target and amplifier of disease^11–13^. Importantly, IL-13-dependent transcriptional programs in EoE are tissue-specific and reflect coordinated interactions between epithelial cells and infiltrating immune cells^14,15^. Although EoG and EoE share clinical and immunological features^16,17^, the extent to which pathogenic mechanisms defined in the esophagus translate to the stomach remains unclear, particularly given their divergent epithelial differentiation, functional specialization, and environmental exposures.

IL-4 and IL-13 are central mediators of type 2 inflammation and exert their biological effects through shared but distinct receptor complexes^18^. IL-4 signals through the type I receptor composed of IL-4Rα and the common γ-chain. IL-4 can also signal via the type II receptor consisting of IL-4Rα and IL-13Rα1. Importantly, the type II IL-4R is shared with IL-13. Thus, IL-4 signals through both type I and type II receptor complexes, whereas IL-13 signals exclusively through the type II receptor^19^. Subsequently, therapeutic targeting of these receptor chains may have overlapping but non-identical consequences in type 2 inflammation^20–22^. Studies in allergic airway disease, atopic dermatitis, and models of food allergy have demonstrated that IL-4Rα signaling is required for IgE class switching, eosinophil recruitment, and mast-cell activation, whereas IL-13Rα1 plays a more specialized role in regulating epithelial responses, barrier dysfunction, and tissue remodeling^12,20,21,23–25^. In contrast, in EoE, IL-13 signaling via IL-13Rα1 is critically required for the full spectrum of histopathological features, including eosinophil infiltration, epithelial hyperplasia, angiogenesis, and fibrosis^12^. These data underscore tissue-specific utilization of these receptor pathways in type 2 inflammation.

Despite advances in our understanding of EoG, the relative contribution of IL-4Rα-and IL-13Rα1-mediated signaling to gastric inflammation and remodeling in EoG remains to be defined. Understanding the roles of these receptor chains is particularly timely given the emergence of biologic therapies targeting type 2 cytokine pathways, including agents directed against IL-4Rα (e.g., dupilumab) and IL-13 (e.g., Lebrikizumab, Cendakimab)^26–28^. Nonetheless, mechanistic dissection of EoG has been limited by the absence of well-characterized experimental models that recapitulate human histopathology including immune-cell accumulation as well as epithelial and fibrotic remodeling.

Herein, we developed a mouse model of experimental EoG induced by cutaneous sensitization followed by repeated intragastric oxazolone challenge. Using this model, we first defined the transcriptional programs associated with gastric type 2 inflammation and remodeling. Subsequently we employed IL-4Rα neutralization and Il13ra1-deficient mice to interrogate receptor-specific functions in immune-cell recruitment, epithelial remodeling, and fibrosis. Our findings identify non-redundant roles for IL-4Rα and IL-13Rα1 in experimental EoG, establishing IL-4Rα as a key mediator of immune and structural pathology while revealing a selective role for IL-13Rα1 in epithelial remodeling. Together, these data provide mechanistic insight into cytokine-receptor signaling in EoG and offer a framework for therapeutic targeting EoG and additional EGIDs.

## Materials and Methods

### Experimental EoG

Mice (male c57BL6 mice, 6-8 weeks) were skin sensitized (15μl 1% OXA, Sigma #E0753, dissolved in acetone) on both ears (60 μl per mouse). After five additional skin challenges (0.5% OXA in acetone), the mice were bled and serum IgE was quantitated (ELISA, BD Bioscience, USA). The mice were intragastrically challenged (day 18, 200µl OXA, 1% in a 1:2 ratio of olive oil and 95% alcohol, respectively) using a plastic feeding tube that reaches the stomach (Instech, 22ga X 25mm). On day 36, the mice were euthanized. A thorough explanation of the methods used in this study and analyses can be found in the supplemental file online.

## Results

### Intragastric oxazolone challenge induces experimental eosinophilic gastritis

To establish an experimental model of EoG, C57BL6/J mice were sensitized epicutaneously on the ear with 1% oxazolone (OXA) on day 0, followed by five additional skin challenges with 0.5% OXA (Figure 1A). Twenty-four hours after the final ear challenge, serum total IgE levels were significantly elevated, confirming systemic type 2 immune activation (Supplementary Figure 1). Mice then underwent eight intragastric challenges with 1% OXA or vehicle control. On day 36, stomachs were collected for histological analysis. OXA-challenged mice exhibited robust eosinophilic infiltration within the gastric mucosa, as demonstrated by anti-major basic protein (MBP) immunostaining (Fig. 1B, C). Chloroacetate esterase (CAE) staining further revealed increased mast-cell accumulation within the gastric mucosa^29^ (Fig. 1D, E).

**Figure 1.**
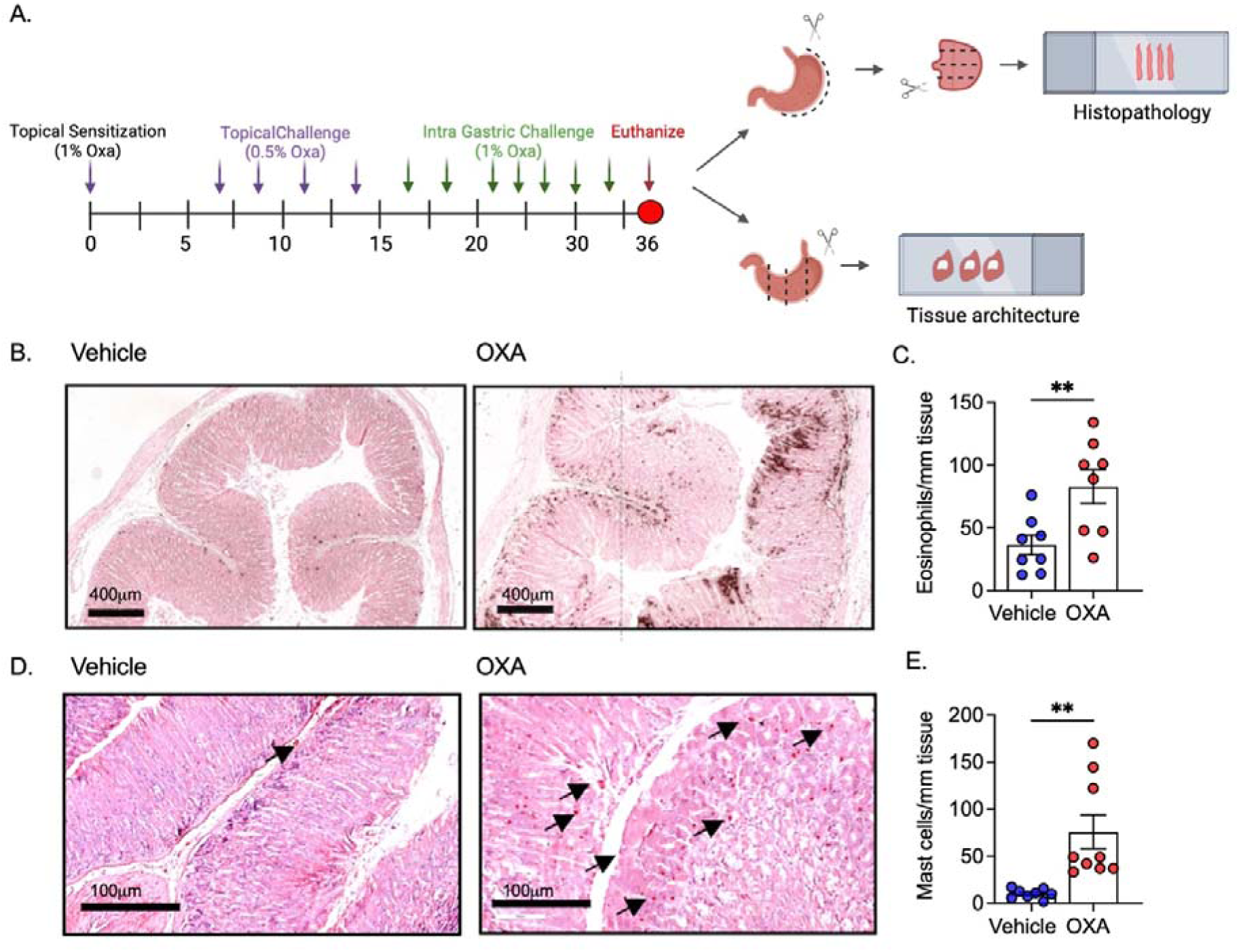
Experimental EoG induces eosinophil and mast-cell infiltration in the gastric mucosa. Schematic illustration of the experimental protocol used to induce eosinophilic gastritis (EoG) by epicutaneous sensitization followed repeated intragastric oxazolone (OXA) challenge (A). gastric sections were collected from vehicle- and OXA-treated mice at day 36 and analyzed for inflammatory cell infiltration (B-E). Eosinophils were detected by anti-major basic protein (MBP) immunostaining (B), and mast cells were identified by chloroacetate esterase (CAE) staining (D). Quantification of MBP^⁺^ eosinophils (C) and CAE^⁺^ mast cells (E) was performed using QuPath software. Data represent mean ± SEM from one representative experiment with n = 8-12 mice per group. In (C) and (E), each dot represents an individual mouse analyzed on day 36. Statistical significance was determined by two-tailed unpaired t test, **- p < 0.01.

### Experimental EoG is characterized by epithelial remodeling and submucosal fibrosis

Histological examination of gastric sections from OXA-challenged mice revealed pronounced structural alterations compared with vehicle-treated controls. Hematoxylin and eosin (H&E) staining demonstrated thickening of the mucosal and submucosal layers and elongation of gastric foveolae (Figure 2A-B). Quantitative analyses confirmed significant increases in mucosal epithelial thickness, submucosal thickness, and foveolar length in OXA-treated mice (Figure 2D-F). Consistent with these changes, Gomori’s trichrome staining showed enhanced collagen deposition within the subepithelial region of OXA-challenged stomachs, indicative of tissue fibrosis (Figure 2C). Quantification of trichrome-positive area revealed a significant increase in subepithelial fibrotic deposition compared with controls (Figure 2G). Taken together, these data demonstrate that experimental EoG is accompanied by key histopathological features of human disease, including epithelial remodeling and subepithelial fibrosis

**Figure 2.**
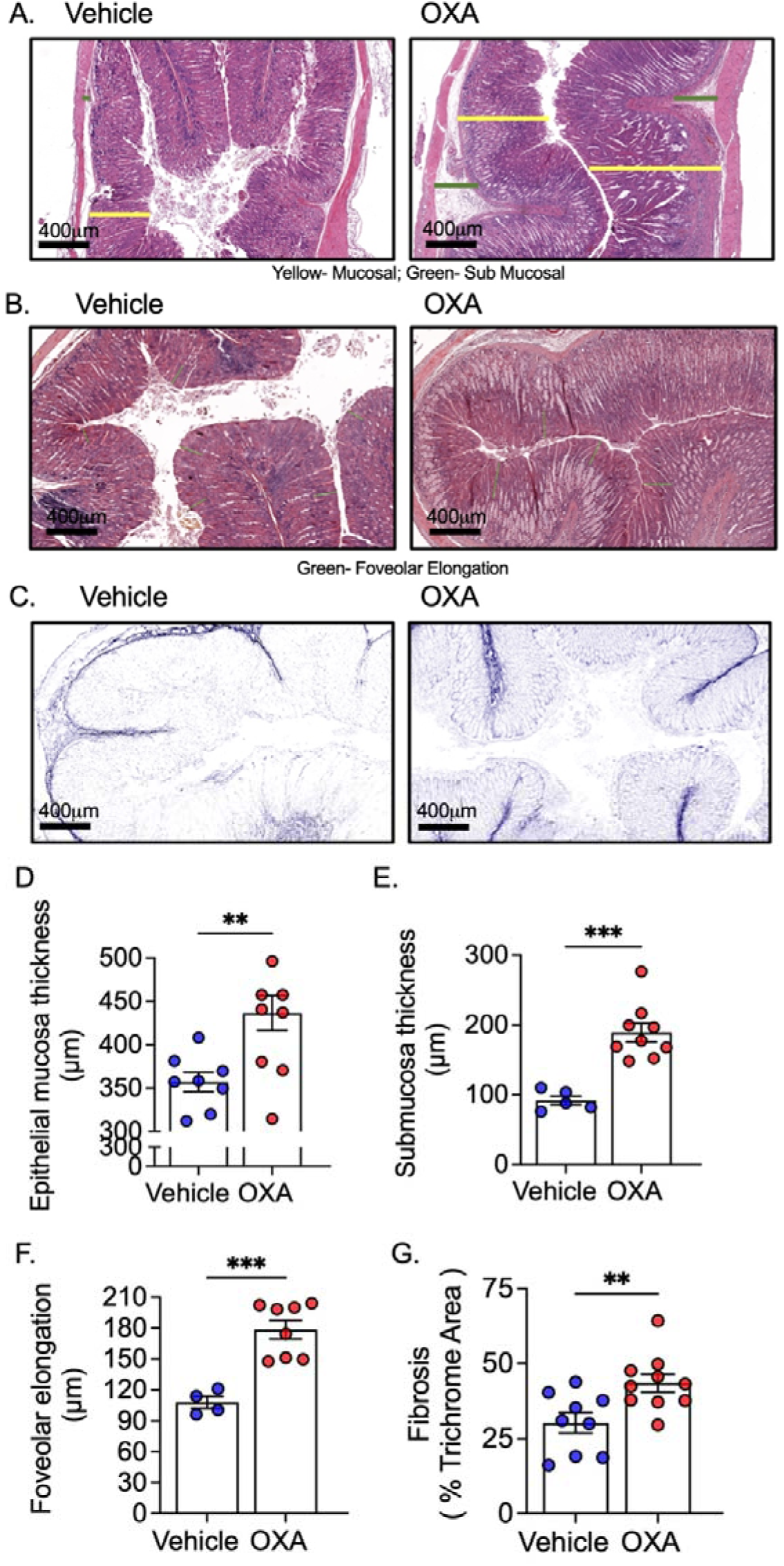
Experimental EoG induces gastric tissue remodeling. Gastric tissue was collected on day 36 from vehicle- and oxazolone (OXA)-treated mice subjected to the experimental EoG protocol. Representative hematoxylin and eosin (H&E) stained gastric sections from vehicle- and OXA-treated mice are shown, illustrating epithelial mucosal and submucosal architecture (A). Quantification of epithelial mucosal thickness and submucosal thickness was performed on H&E-stained sections (D, E). Additional representative H&E-stained gastric sections show foveolar architecture in vehicle- and OXA-treated mice (B). Quantification of foveolar elongation was performed from H&E-stained sections (F). Representative Gomori’s Trichrome-stained gastric sections from vehicle- and OXA-treated mice are shown (C). Quantification of Trichrome-positive area (% area) was used to assess collagen deposition (G). Data represent mean ± SEM from a representative experiment with n = 8-12 mice per group. Statistical significance was determined by unpaired two-tailed t test, **- p < 0.01, ***- p <0.001.

### Experimental EoG induces a transcriptional program charactarized by IL-4/IL-13-associated epithelial remodeling and fibrosis

To define the molecular programs associated with experimental EoG, bulk RNA sequencing was performed on gastric tissue obtained from vehicle- and OXA-challenged mice collected on day 36. Global transcriptomic analysis revealed marked differences between OXA-challenged and vehicle-treated stomachs. Principal component analysis demonstrated clear separation of samples along the first principal component, which accounted for 74% of the total variance, indicating a robust and coordinated transcriptional response to intragastric OXA challenge (Figure 3A). Consistent with this separation, hierarchical clustering identified extensive transcriptomic reorganization in OXA-treated stomachs relative to controls, with 2872 genes differentially expressed (p<0.05, fold change ≥1.5), including 1530 upregulated and 1374 downregulated transcripts (Figure 3B and Table S1-3).

**Figure 3.**
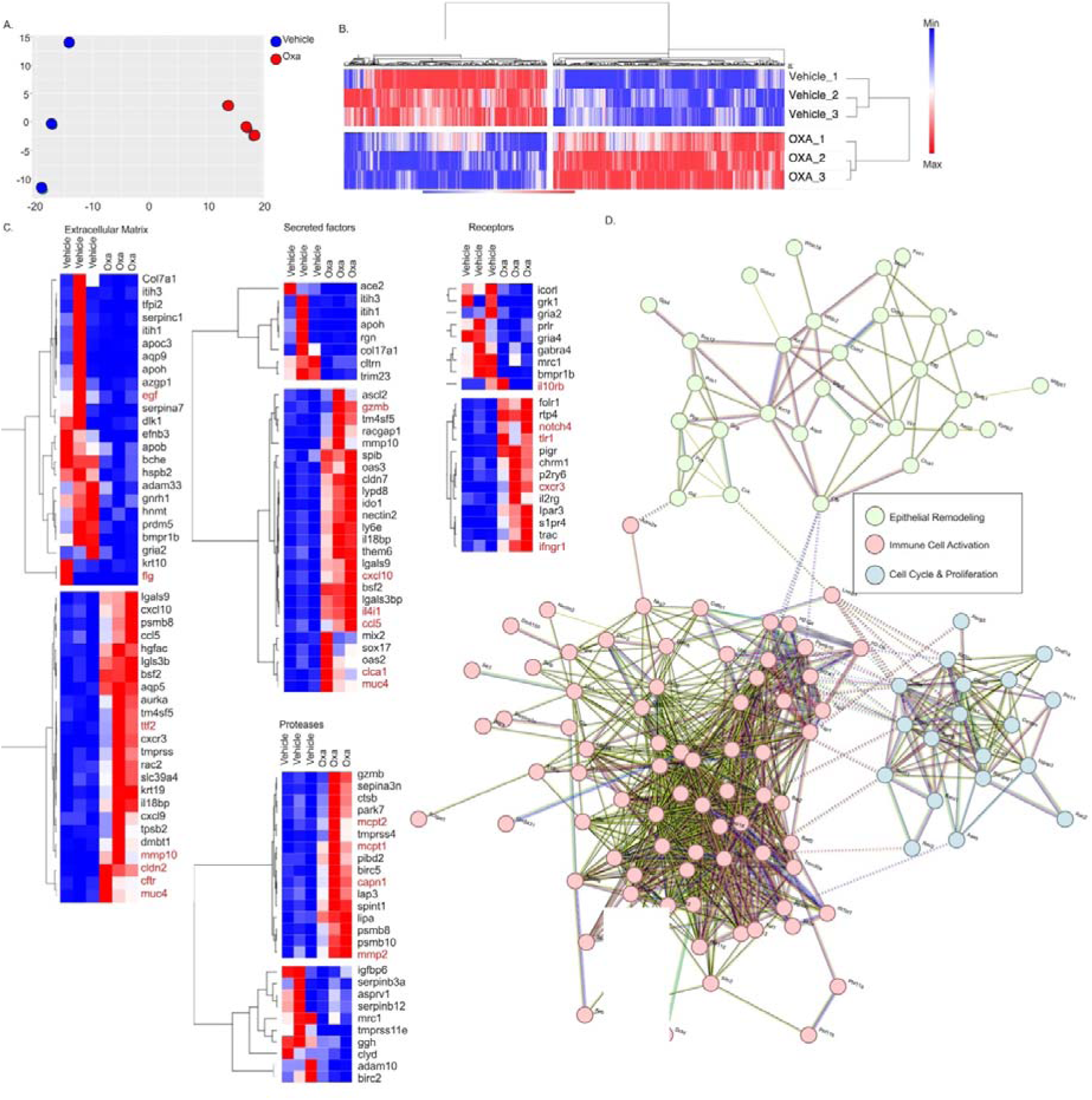
RNA sequencing of experimental EoG identifies transcriptional programs associated with epithelial remodeling and immune activation. Experimental eosinophilic gastritis (EoG) was induced in wild-type mice, and gastric tissue was collected on day 36 from vehicle- and oxazolone (OXA)-treated animals and subjected to bulk RNA sequencing analysis. Principal component analysis (PCA) demonstrates separation of transcriptomic profiles between vehicle- and OXA-treated samples along the first principal component (A). A heat map of differentially expressed genes illustrates global transcriptional changes between vehicle- and OXA-treated stomachs (B). Heat maps display of selected differentially expressed genes grouped by functional categories, including extracellular matrix-associated genes, secreted factors, and proteases as well as receptor-encoding transcripts (C). Expression values are shown for individual samples from vehicle- and OXA-treated mice. STRING network analysis of upregulated genes identifies interconnected gene modules corresponding to epithelial remodeling, immune cell activation, and cell cycle/proliferation programs (D). Nodes are colored according to functional classification, as indicated. Data shown are representative of one experiment, with each column corresponding to an individual mouse (n=3 per group)

Analysis of upregulated genes revealed a dominant transcriptional signature associated with epithelial activation, tissue remodeling, and inflammatory responses. Among epithelial-derived transcripts, multiple genes implicated in epithelial repair and barrier modulation were robustly induced, including *Areg, Tff2, Tff1, Muc4, Muc1, Muc6, Cldn2, and Clca1* (Figure 3C). In parallel, genes encoding extracellular matrix components and fibrosis-associated factors were upregulated, including multiple collagen genes (*Col1a1, Col4a1, Col4a2, Col6a1, Col6a2, Col6a4, Col15a1, Col18a1*), as well as matrix-associated regulators such as *Tgfbi* and *Loxl2* (Table S2 and S4).

Genes encoding proteases and mast-cell effector molecules were also prominently induced, including *Mmp10, Mmp15, Mmp2, Mcpt1, Mcpt2, Tpsb2, Cma1*, and *Cpa3*, consistent with the increased mast-cell accumulation observed histologically (Figure 1 and Tables S2 and S4). Upregulation of cytokine-responsive and signaling components, including *Il4ra* and *Il4i1*, further indicated engagement of IL-4/IL-13-associated pathways in the gastric mucosa.

STRING network analysis of upregulated genes identified several interconnected modules, including three main clusters. An epithelial remodeling and junctional cluster, a proliferation-associated cluster, and an immune activation cluster enriched for antigen processing and immune cell activation-associated genes (Figure 3D).

In contrast to the induction of remodeling and inflammatory programs, a distinct subset of genes was significantly downregulated in OXA-challenged stomachs, reflecting suppression of steady-state gastric epithelial and homeostatic functions. Multiple transcripts encoding ion transporters and channels were reduced, including *Slc26a7, Slc12a2, Slc16a7, Slc27a2, Kcne2*, and *Kcnj16*. Genes involved in gastric acid secretion and epithelial maintenance, including *Atp4a* and *Egf*, were also markedly downregulated (Figure 3C, Table S3 and S4).

Pathway enrichment analysis of downregulated genes revealed significant overrepresentation of biological processes related to ion transport, transmembrane transport, metabolic processes, and chemical homeostasis. Molecular function analysis highlighted transporter activity, ion channel activity, and ATPase activity, while cellular component analysis identified plasma membrane and mitochondrial compartments as major sites of suppression (Figure 4A-B). STRING network analysis of downregulated genes demonstrated a highly interconnected network dominated by metabolic detoxification, and transport machinery, including cytochrome P450 enzymes, phase I/II metabolic genes, and epithelial transport components, indicating coordinated suppression of differentiated gastric epithelial functional programs (Figure 4C).

**Figure 4.**
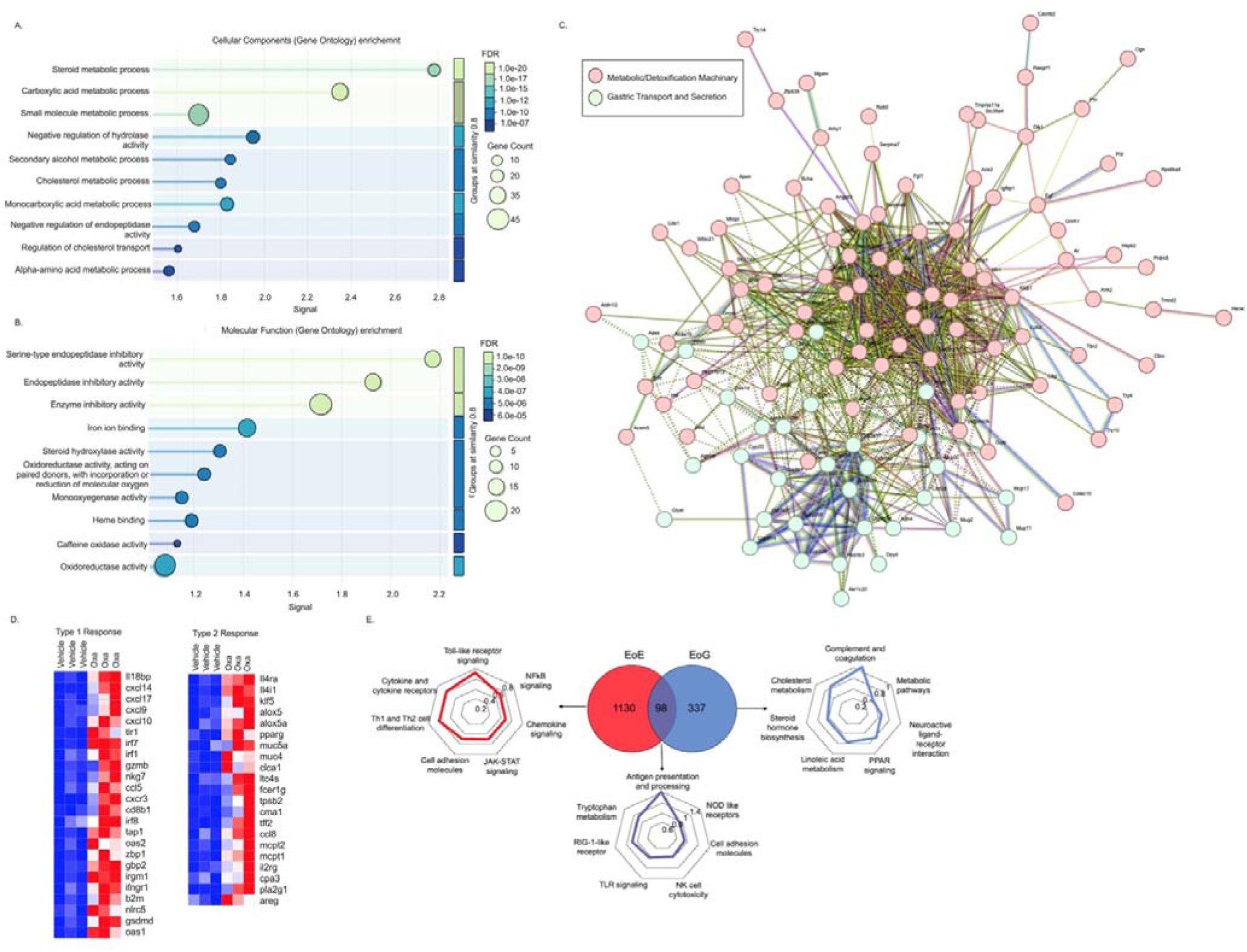
Downregulated epithelial gene programs and immune transcriptional signatures in experimental EoG. Bulk RNA sequencing was performed on gastric tissue collected on day 36 from vehicle- and oxazolone (OXA)-treated mice. Differentially expressed genes were analyzed to characterize transcriptional changes associated with experimental eosinophilic gastritis (EoG). Gene Ontology enrichment analysis of biological processes associated with genes downregulated in OXA-treated stomachs is shown (A). Gene Ontology enrichment analysis of molecular functions associated with downregulated genes is shown (B). In (A) and (B), dot size represents the number of genes associated with each term, and color indicates false discovery rate (FDR). STRING network analysis of downregulated genes is shown, illustrating protein-protein interaction networks among genes suppressed in experimental EoG (C). Heat maps display expression of selected genes associated with type 1 (left) and type 2 (right) immune responses in vehicle- and OXA-treated samples (D), with each column representing an individual mouse. Venn diagrams and radar plots compare differentially expressed genes in experimental EoG with published eosinophilic esophagitis (EoE) datasets in mouse (E).

### Experimental EoG exhibits a mixed immune transcriptional profile and suppression of gastric homeostatic programs distinct from EoE

Further examination of immune-associated transcripts demonstrated that experimental EoG is characterized by a composite inflammatory signature comprising both type 2- and interferon-associated components (Figure 4D). Type 2-associated genes, including *Il4ra, Il4i1, Mcpt1, Mcpt2, Muc4, Areg, and Tff2*, were induced (Table S5). In parallel, multiple genes associated with interferon signaling and cytotoxic immune responses were also elevated (Table S5), including *Ifngr1, Irf1, Irf7, Irf8, Tap1, Cxcl10, Cxcl9, Cd8b, and Gzmb*, indicating concurrent activation of type 2 and interferon-associated immune pathways.

To place the experimental EoG transcriptome in the context of other EGIDs, we compared the experimental EoG gene signature with published experimental EoE datasets^12^. This analysis revealed that a large fraction of the differentially expressed genes was disease-specific, with 1,130 transcripts uniquely enriched in EoE and 337 transcripts uniquely enriched in EoG, whereas a smaller but coherent set of 98 genes was shared between the two conditions (Figure 4E).

The EoE-specific gene signature (Table S6) was enriched for pathways related to cytokine and chemokine signaling, NFκB and JAK-STAT signaling, Toll-like receptor pathways, and cell adhesion, consistent with an immune communication and recruitment phenotype. Representative EoE-unique genes included chemokines and chemokine receptors (*Ccl1, Ccl3, Ccl4, Ccl8, Ccl22, Cxcl11, Ccr7*), T cell-associated signaling molecules (*Cd3d, Cd3e, Cd3g, Lck, Zap70, Lat, Itk*), immune regulatory receptors (*Pdcd1, Lag3, Havcr2, Cd274*), and squamous epithelial differentiation, barrier-associated genes such as keratins (*Krt5, Krt14, Krt16, Krt17*) and cornified envelope components (*Flg, Sprr, Lce*).

In contrast, the EoG-specific gene signature (Table S7) was enriched for pathways associated with metabolic processes, lipid and cholesterol metabolism, complement, coagulation cascades, PPAR signaling, and neuroactive ligand-receptor interactions (Figure 4E). EoG-unique genes included apolipoproteins and lipid-handling factors (*Apob, Apoa2, Apoc1/3/4, Angptl3, Mttp*), metabolic and detoxification enzymes (*Cyp1a2, Cyp3a11, Cyp2c family members, Mat1a, Gnmt, Bhmt*), gastric epithelial transport and interface genes (*Slc26a7, Cftr, Ace2, Pigr*), together with remodeling-associated transcripts (*Muc4, Mmp10*) and mast cell-associated genes (*Tpsb2, Ms4a2*). This signature is consistent with glandular epithelial remodeling and altered gastric homeostatic programs.

The shared EoE-EoG gene set (Table S8), defined a conserved inflammatory module enriched for antigen processing and presentation, interferon-associated signaling, and cytotoxic immune programs. Shared genes included antigen-presentation machinery (*B2m, Tap1, Tap2, Tapbp, H2-K1, H2-D1*), interferon-stimulated genes (*Irf7, Mx1, Isg15, Oas family members, Dhx58, Ddx58*), and cytotoxic lymphocyte markers (*Nkg7, Gzmb, Cd8b1, Trac, Ccl5*). Notably, this shared signature also contained a type 2-associated effector and epithelial remodeling component, exemplified by mast cell proteases (*Mcpt1, Mcpt2*), the immunoregulatory enzyme *Il4i1*, and epithelial stress-response genes (*Clca1, Tff2*).

### IL-4R**α** signaling regulates immune-cell infiltration and epithelial remodeling in experimental EoG

To determine the functional contribution of IL-4Rα signaling to experimental EoG, mice were treated with an IL-4Rα-neutralizing antibody during the intragastric OXA challenge phase. Histological analysis revealed that anti-IL-4Rα treatment markedly reduced eosinophil and mast-cell accumulation within the gastric mucosa (Figure 5A, 5C). Quantitative analysis confirmed significant decreases in MBP^⁺^ eosinophils and CAE^⁺^ mast cells in the stomachs of anti-IL-4Rα-treated mice compared with isotype-treated controls (Figure 5B, 5D).

**Figure 5.**
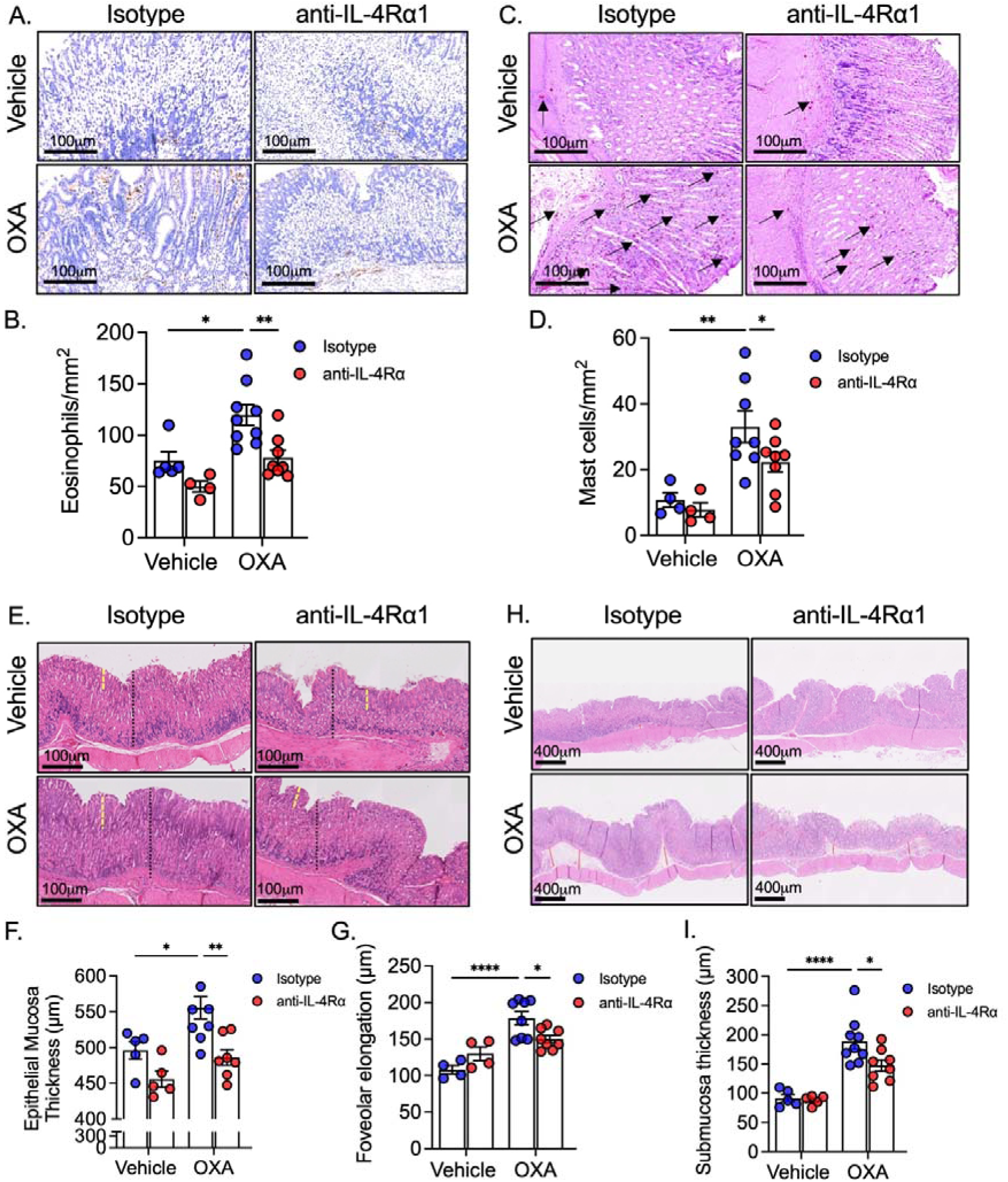
IL-4Rα regulates inflammatory cell infiltration and tissue remodeling in experimental EoG. Experimental EoG was induced in wild-type mice by skin sensitization with oxazolone, followed by administration of isotype control or anti-IL-4Rα antibody during the intragastric challenge phase. Representative gastric sections stained for major basic protein (MBP) are shown to visualize eosinophil infiltration across treatment groups (A), with corresponding quantification of MBP^⁺^ eosinophils (B). Chloroacetate esterase (CAE) staining was used to identify mast cells in gastric sections (C), and mast-cell numbers were quantified (D). Hematoxylin and eosin (H&E)-stained gastric sections illustrate mucosal architecture, including epithelial thickness, foveolar length, and submucosal structure across treatment groups (E, H). Quantitative analysis of foveolar elongation (F), epithelial mucosal thickness (G), and submucosal thickness (I) was performed on H&E-stained sections. Data represent mean ± SEM from a representative experiment using n = 8-12 mice per group. Statistical significance was determined using one way ANOVA followed by Tukey-Kramer post-hoc test; *-p<0.05, **- p < 0.01, ***- p <0.001.

In addition to its effects on immune-cell infiltration, IL-4Rα blockade substantially attenuated epithelial remodeling. Hematoxylin and eosin-stained sections demonstrated marked improvement in mucosal architecture, characterized by reduced epithelial thickening, shortened foveolar length, and diminished submucosal expansion (Figure 5E, 5H). Histological measurements confirmed significant reductions in epithelial mucosal thickness, foveolar elongation, and submucosal thickness in anti-IL-4Rα-treated mice relative to isotype controls (Figure 5F, G, and I).

Together, these findings identify IL-4Rα as a central regulator of both immune-cell infiltration and structural remodeling in experimental EoG.

### IL-13R**α**1 selectively regulates epithelial remodeling but not immune-cell infiltration in experimental EoG

To determine the role of IL-13Rα1 signaling in experimental EoG, *Il13ra1^-/-^* mice and wild-type (WT) controls were subjected to the oxazolone-induced EoG protocol. Deficiency in *Il13ra1* did not affect inflammatory cell infiltration. Immunostaining for MBP revealed comparable numbers of eosinophils in OXA-treated WT and *Il13ra1^-/-^*mice (Figure 6A-B). Similarly, chloroacetate esterase (CAE) staining demonstrated equivalent mast-cell densities between genotypes following OXA challenge (Figure 6C-D). In contrast to its effects on eosinophil and mast cell accumulation, histological analysis revealed that loss of *Il13ra1* markedly attenuated epithelial remodeling in response to OXA challenge (Figure 6E-H). Hematoxylin and eosin-stained sections from *Il13ra1^-/-^* mice exhibited reduced epithelial thickening and shortened foveolar projections compared with WT controls (Figure 6E). Quantitative morphometric analysis confirmed significant reductions in epithelial mucosal thickness and foveolar elongation in *Il13ra1^-/-^* mice (Figure 6F-H).

**Figure 6.**
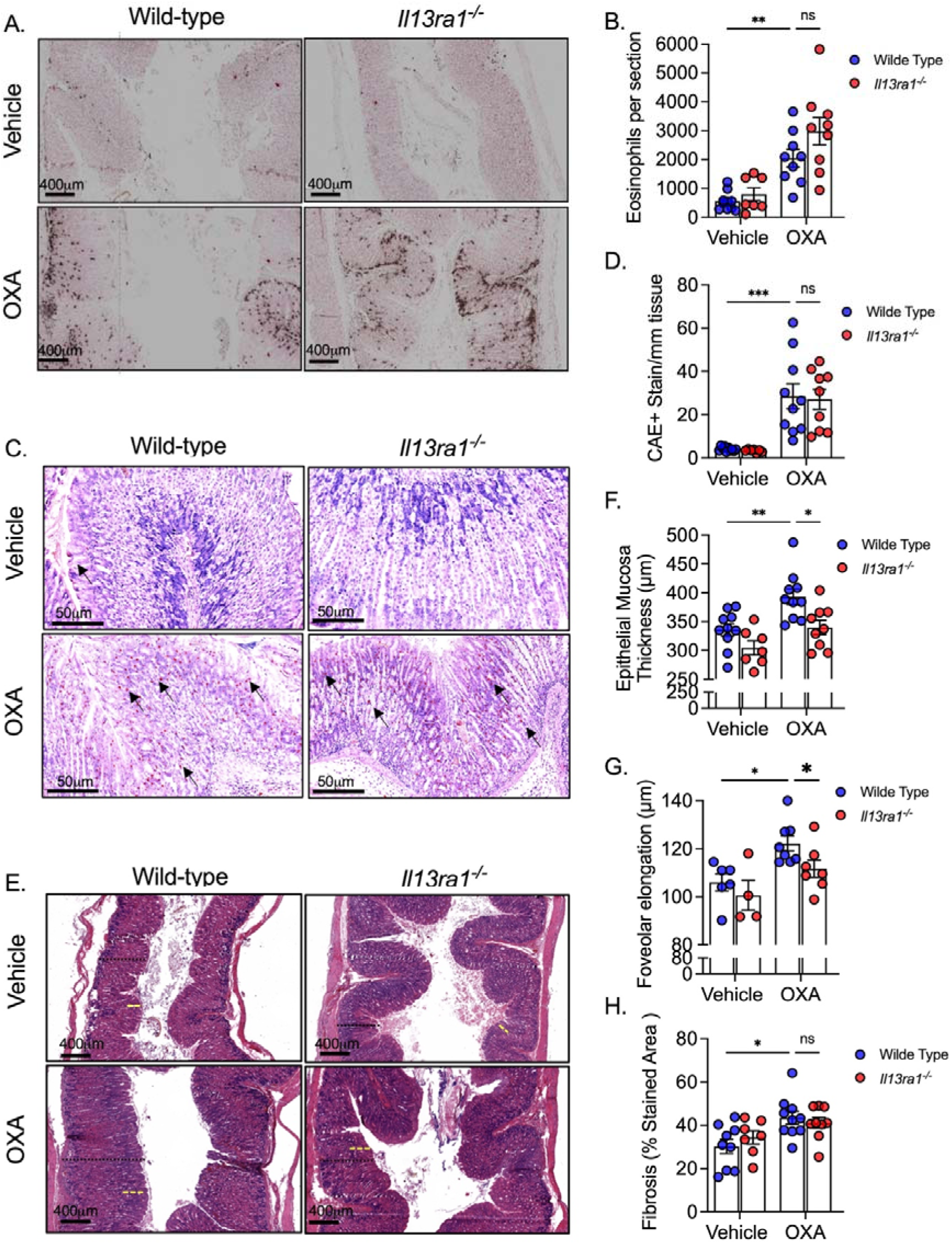
IL-13Rα1 regulates tissue remodeling but not inflammatory cell infiltration in experimental EoG. Experimental EoG was induced in wild-type and *Il13ra1^-/-^* mice. On day 36, gastric tissue was collected, processed and stained for major basic protein (MBP) are shown (A), with corresponding quantification of total MBP^⁺^ eosinophils per gastric section (B). Representative chloroacetate esterase (CAE) stained gastric sections are shown to visualize mast cells (C), and CAE^⁺^ mast cells per mm² of tissue were quantified (D). Representative hematoxylin and eosin (H&E)-stained gastric sections from vehicle- and OXA-treated wild-type and *Il13ra1^-/-^* mice are shown, illustrating epithelial mucosal architecture (E). Quantification of epithelial mucosal thickness and foveolar elongation was performed on H&E-stained sections (F, G). Representative gastric sections stained. Data represent mean ± SEM from a representative experiment using n = 8-12 mice per group. Statistical significance was determined by two-way ANOVA followed by Tukey–Kramer post hoc test; **-p < 0.01, ***-p < 0.001; ns, not significant).

Together, these findings identify IL-13Rα1 as a selective regulator of epithelial remodeling in experimental EoG, acting independently of the IL-4Rα-dependent pathways that govern eosinophil and mast-cell accumulation.

## Discussion

EoG is a poorly understood EGID, and the cytokine pathways that coordinate immune-cell infiltration and tissue remodeling in the stomach remain to be defined. Progress in understanding these mechanisms has been limited by restricted access to human tissue and by the lack of experimental models that recapitulate both inflammatory infiltration and epithelial remodeling in the gastric mucosa. IL-4 and IL-13 are hallmark cytokines of type 2 immune responses and represent central targets for therapeutic intervention^30^. They mediate their biological effects through overlapping but non-identical receptor complexes that are differentially expressed across immune and non-immune compartments^31^, suggesting that tissue pathology may be shaped in a receptor-dependent manner. Notably, although IL-4 and IL-13 have been implicated in EoG^1,7–9,32^, the relative contribution of their shared and distinct receptor pathways in the stomach has not been determined. In this study, we established a robust experimental model of EoG and used it to dissect the receptor-specific contributions of IL-4Rα and IL-13Rα1 to gastric type 2 inflammation. Our data demonstrate that IL-4Rα signaling is required for both eosinophil and mast-cell accumulation and for epithelial and subepithelial remodeling, whereas IL-13Rα1 selectively regulates epithelial remodeling without affecting inflammatory cell infiltration. These findings identify non-redundant roles for IL-4/IL-13 receptor pathways in EoG and provide a mechanistic framework for understanding gastric eosinophilic disease.

The experimental EoG model described here reproduces key pathological and molecular features that have been reported in human disease^3^. Histologically, intragastric OXA challenge induced dense eosinophil and mast-cell infiltration of the gastric mucosa together with epithelial hyperplasia, foveolar elongation, and subepithelial fibrosis^33,34^. These features closely mirror those observed in gastric biopsies from patients with EoG^3^. Importantly, human studies have emphasized that epithelial remodeling, periglandular fibroplasia, and smooth muscle hyperplasia are central components of EoG pathology and are not simply secondary to eosinophil accumulation^35,36^. The presence of these structural changes in experimental EoG supports its relevance for mechanistic interrogation of gastric eosinophilic disease.

At the molecular level, the histologic features which were observed were accompanied by a transcriptional program dominated by epithelial activation and tissue remodeling. For instance, genes associated with epithelial repair and barrier modulation, including *Areg*, *Tff2*, *Muc4*, *Cldn2*, and *Clca1*, were upregulated^37–39^. In addition, proteases and matrix-associated genes such as *Mmp10*, *Mcpt1*, *Mcpt2*, as well as multiple collagens were upregulated as well. This pattern aligns with transcriptomic analyses of human EoG biopsies that identified a conserved gastric gene signature enriched for epithelial-derived factors, extracellular matrix remodeling, and eosinophil-associated pathways, (e.g., CCL26, IL13RA2, MUC4, and AREG)^36^. In contrast, and consistent with epithelial injury and loss of homeostatic gastric programs in EoG, several genes associated with normal gastric epithelial function and ion transport were similarly downregulated in our experimental system and in human EoG (e.g., *Slc26a7* and *Atp4a*)^36^. Notably, both human and experimental EoG engage type 2 inflammatory pathways similar to those described in eosinophilic esophagitis (EoE)^6,40^. However, comparative transcriptomic analyses demonstrate that only a minority of dysregulated genes are shared between the two diseases, indicating that common inflammatory cues are interpreted in a tissue-specific manner. The shared EoE-EoG gene set defines a conserved inflammatory module enriched for antigen processing and presentation, interferon-associated signaling, cytotoxic lymphocyte markers, and a type 2 effector and epithelial stress-response component. In contrast, disease-specific gene programs reflect the distinct epithelial architectures and physiological functions of the esophagus and stomach, with EoE characterized by squamous epithelial differentiation and immune recruitment pathways, and EoG dominated by glandular metabolic, transport, and remodeling programs. Together, these findings support a model in which EGIDs share a core inflammatory backbone, upon which tissue-specific transcriptional programs are layered to generate disease-defining pathology.

Interestingly, the shared EoE-EoG inflammatory module was strongly enriched for interferon-associated genes, including interferon-stimulated genes, antigen-processing machinery, and IFN-γ-responsive chemokines. Although EoE and EoG are classically viewed as type 2-driven diseases^4,41,42^, accumulating evidence indicates that IFN-γ signaling contributes to disease pathogenesis in EoE^43–45^. Our findings extend these observations by demonstrating that IFN-γ-associated pathways represent a conserved component of experimental EGIDs, coexisting with type 2 effector programs.

Using antibody-mediated neutralization of IL-4Rα and genetic deletion of *Il13ra1*, we identified distinct roles for IL-4Rα and IL-13Rα1 in experimental EoG that are consistent with the organization of type I and type II IL-4 receptor signaling in immune and non-immune cellular compartments. Blockade of IL-4Rα markedly reduced eosinophil and mast-cell accumulation in the gastric mucosa and substantially attenuated epithelial thickening, foveolar elongation, and subepithelial remodeling. These findings indicate that IL-4Rα-dependent signaling is required upstream of both inflammatory cell recruitment and tissue remodeling in the stomach. Because IL-4Rα is shared by the type I receptor, which is predominantly expressed by hematopoietic cells, and the type II receptor, which is broadly expressed by epithelial and stromal cells, IL-4Rα blockade likely disrupts signaling in both immune and non-immune compartments^19,31^. In contrast, genetic deletion of *Il13ra1* selectively affected epithelial remodeling without altering eosinophil or mast-cell accumulation. This dissociation indicates that IL-13Rα1-dependent signaling is dispensable for inflammatory cell recruitment but is required for execution of epithelial remodeling programs. The persistence of eosinophilic and mast-cell infiltration in *Il13ra1^-/-^* mice suggests that inflammatory recruitment in EoG is driven primarily through IL-4 via the type I IL-4 that does not require IL-13 and the type II IL-4R.

This functional separation aligns with observations in other type 2 diseases. In models of allergic airway disease and atopic dermatitis, IL-13Rα1-dependent signaling predominantly regulates epithelial differentiation, mucus-associated gene expression, and tissue remodeling, whereas interventions targeting IL-4Rα exert broader effects that include suppression of inflammatory cell accumulation^20,21,23,24^. The pattern observed in experimental EoG therefore extends this paradigm to the gastric mucosa. Importantly, this organization differs from EoE, in which IL-13Rα1 signaling has been linked to both epithelial remodeling and inflammatory features^12^. Given the frequent coexistence of EoE and EoG in patients^46–49^, our findings suggest that IL-4/IL-13 receptor pathways are deployed in a tissue-specific manner across the gastrointestinal tract, with distinct consequences for immune recruitment and structural remodeling.

The receptor-specific organization of type 2 cytokine signaling identified here has direct translational implications for therapeutic strategies in EoG. Clinical experience with dupilumab, which targets IL-4Rα and thereby inhibits signaling by IL-4 and IL-13, has demonstrated histologic and symptomatic benefit in EoE and is now being explored in non-esophageal EGIDs, including EoG ^1,7,9,50,51^. Although published data in EoG remain limited to case reports, small series, and early-phase trials, these studies consistently suggest that IL-4Rα blockade can reduce tissue eosinophilia and improve disease activity. Our findings provide a mechanistic framework for these observations. Based on our data, selective disruption of IL-13 signaling is predicted, to preferentially affect epithelial remodeling while having less impact on eosinophil and mast-cell accumulation. This distinction may be clinically relevant, particularly in patients with EoG who exhibit prominent structural changes such as epithelial hyperplasia, foveolar elongation, or fibrosis. More broadly, these findings suggest that differential targeting of IL-4Rα versus IL-13Rα1-dependent pathways could yield distinct therapeutic effects across coexisting EGID manifestations. Since EoG frequently occurs alongside EoE^46–49^, selective interventions may have divergent effects across gastrointestinal segments. This underscores the need to consider tissue-specific cytokine signaling when designing and interpreting therapeutic strategies.

This study has several limitations. First, our findings are based on a murine model of experimental EoG, which, may not reflect the full heterogeneity or chronicity of EoG in patients. Second, transcriptomic analyses were performed on bulk gastric tissue, which limits cell-type specific resolution and precludes direct attribution of gene expression changes to individual cellular populations. Finally, although our findings align with published human EoG transcriptomic studies and emerging clinical experience with IL-4Rα-targeted therapies, direct validation in human gastric tissue was not performed. Future studies integrating single-cell approaches, spatial analyses, and human biopsy validation will be important to further refine the receptor-specific mechanisms identified here.

In summary, we establish a robust experimental model of EoG and define receptor-specific roles for IL-4Rα and IL-13Rα1 in gastric type 2 inflammation. These findings provide mechanistic insight into how IL-4 and IL-13 signaling pathways are differentially deployed in the gastric mucosa and highlight EoG as a biologically distinct EGID shaped by tissue-specific interpretation of shared cytokine signals. Furthermore, the study provides important and clinically relevant insights regarding the molecular pathways mediating gastric eosinophilic disease.

## Acknowledgments

We wish to thank all the members of the Munitz lab for their constructive comments.

## Author Contributions

These authors contributed as follows:

AD and AM- conception and/or design of the work

AD, SS, SB and MI - Data Collection

AD, SS, SB, MI and AM- Data analysis and interpretation.

AD, and AM- Drafting the article

AD, SS, SB, MI and AM - Critical revision of the article

AM- Final approval of the version to be published

