## Supplementary Text for "Differential roles for IL-4Rα and IL-13Rα1 in immune cell infiltration and epithelial remodeling in experimental eosinophilic gastritis"

^2^ The Roberts-Guthman Chair in Immunopharmacology

**Short Title:** IL-4 and IL-13 receptors in eosinophilic gastritis

**Word count:**

**Key words:** EGIDs, Eosinophilic Gastritis, IL-4, IL-13, IL-13Rα1, IL-4Rα

**^*^Corresponding author:** Ariel Munitz, PhD, The Roberts-Guthman Chair in Immunopharmacology, Department of Clinical Microbiology and Immunology, Gray Faculty of Medical and Health Sciences, Tel Aviv University, Ramat Aviv 69978, Israel. Tel. (Office): +972-3-640-7636, Fax: +972-3-640-9160,; ORCID: <https://orcid.org/0000-0003-1626-3019>;

**Materials and methods:**

**Mice**

Wild-type (WT) C57BL/6 mice were obtained from Harlan Laboratories (Rehovot, Israel) and grown in-house*. Il13ra1^-/-^* mice^1^, were grown in house. In all experiments, age-, weight-, and sex-matched mice were used and housed under specific-pathogen-free conditions. All experiments were reviewed and approved by the Animal Care Committee of Tel Aviv University and were performed in accordance with its regulations and guidelines regarding the care and use of animals for experimental procedures.

**Experimental eosinophilic gastritis (EoG)**

Experimental EoG was induced as previously described^2^, with slight modifications. Male c57BL6 mice (6–8 weeks) were skin sensitized with 15 μL of 1% Oxazolone (Sigma, #E0753) solution dissolved in acetone on both ear flanks (60 μL per mouse). After five additional skin challenges (0.5% OXA in acetone), the mice were bled and serum IgE was quantitated by ELISA (BD Bioscience, #555248). On Day 18, the mice were intragastric challenges (8 challenges) with 200 μL OXA (1% in a 1:2 ratio of olive oil and 95% alcohol, respectively) using a plastic feeding tube (Instech, #FTP-22-25). The mice were euthanized on day 36.

**In vivo neutralization experiments**

On Day 19 of the experimental EoE protocol, mice were intraperitoneally injected with 150 µg of anti-mouse IL-4Rα neutralizing antibody (clone M1, kindly provided by Dr. Fred Finkelman, Cincinnati Children’s Hospital Medical Center) or 150 µg isotype control rat IgG2a (BioXcell). Additional injections were given twice a week until day 36. On day 36 ,the mice were euthanized, and samples were taken for analysis.

**Histology and immunohistochemistry**

Gastric tissue was harvested, fixed in 4% formaldehyde (Bio-Lab Ltd., Israel), and paraffin-embedded. Sections (5 µm) were deparaffinized and stained with hematoxylin and eosin (H&E; Sigma, Israel). Immunohistochemistry was performed using anti–major basic protein (MBP; kindly provided by Dr. Elizabeth A. Jacobsen, Mayo Clinic, Scottsdale, AZ) and anti–Ki-67 (Novus, #NB110-89717SS). Chloroacetate esterase (CAE) staining was performed on paraffin sections following deparaffinization and incubation at room temperature with a freshly prepared naphthol AS-D chloroacetate solution (Sigma, N0758) containing hexazotized New Fuchsin (New Fuchsin; Fisher Scientific, AC21214; sodium nitrite; Sigma, 237213), followed by hematoxylin counterstaining, dehydration, and mounting. Masson’s trichrome staining was performed using a Leica multistainer with the Leica Trichrome Staining Kit (3804010802) according to the manufacturer’s instructions. Slides were scanned using an Aperio slide scanner (Leica) and analyzed using ImageScope (Leica) and ImageJ.

**RNA sequencing**

The stomach was dissected, and RNA extracted using TRIzol reagent (Invitrogen, #15596026). RNA was quantified using a Qubit Flex Fluorometer (Invitrogen) with the Equalbit RNA High Sensitivity Assay Kit (Vazyme, #EQ211). CEL-Seq2 libraries were prepared as described^3^, with modifications to use 2 ng of purified RNA as input instead of single cells. Sequencing was performed on an Illumina NextSeq2000 platform using P2 100 cycles (Read1-12; Index1-6; Index2-0; Read2-65). Reads were demultiplexed following the CEL-Seq2 pipeline with parameters: min_bc_quality = 10, bc_length = 6, umi_length = 6, and cut_length = 70. Quality control was performed using FastQC (v0.11.9), and reads were trimmed for adapters and poly-A tails using CUTADAPT (v4.4) with a minimum Phred score of 20 and a minimum length of 25 bp. Reads were aligned to the Mus musculus GRCm39 genome using HISAT2 (v2.2.1) with up to two mismatches allowed per read, and only uniquely mapped reads were retained using SAMtools (v1.13). Gene-level read counts were obtained using HTSeq (v2.0.4) in 'union' mode, and normalization and differential expression analysis were conducted using DESeq2 (v1.28.0). Sample preparation and sequencing were conducted by the "Technion Genome Center”, Life Science and Engineering Interdisciplinary Research Center, Technion, Haifa, Israel.

**Bioinformatics analysis**

The differential expression (DE) of the sequenced genes was calculated using the DEseq2 package in R language on RStudio. Genes were regarded as differentially expressed if they presented a false discovery rate (FDR) adjusted P-value below 0.05, and $log2(fold change)\pm0.58$. The samples were presented using principal component analysis (PCA) created with the ggplot R package. Volcano plots (that visualize results of differential expression analyses) were created using the Enhanced Volcano package in R.

Expression heat maps of selected genes of interest (GOIs) were generated using the Morpheus software (Broad Institute). The expression data was subsetted from the sequencing dataset and scaled per gene (row) to enable direct comparison across genes. Hierarchical clustering was performed using the built-in clustering method in Morpheus. The heatmaps visually represent the relative expression levels of GOIs across experimental conditions or samples.

To explore protein-protein interactions (PPIs) of selected GOIs, analyses were conducted using the STRING database. Genes were queried to identify known and predicted interactions based on experimental evidence, computational predictions, and curated databases. The STRING output was used to construct interaction networks and radar plots aiding the identification of functional pathways and complexes potentially involved in the observed experimental outcomes.

**Statistical analysis**

*P* values of mouse data sets were determined by one-way analysis of variance (ANOVA), unpaired two-tailed Student’s *t-test* with a 95% confidence interval. All statistical tests were performed with GraphPad Prism V10 software. Data are shown as mean ± SEM. *-p < 0.05; ***-p* < 0.01; ****-p* < 0.001.

**Supplementary Figure 1**

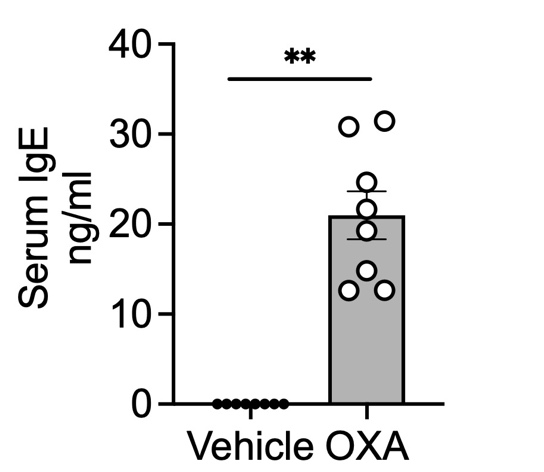

**Figure legend.**

**Figure 1. Experimental EoG is associated with increased serum IgE following skin sensitization.**

To establish an experimental model of EoG C57BL6 mice were skin-sensitized [day 0, 1% oxazolone (OXA)]. Subsequently, the mice received five additional skin-challenges (0.5% OXA, Figure 1A). Twenty-four hours after the last challenge, serum was collected and IgE levels determined. Data are representative of one out of three experiments, each circle represents one mouse, **-p<0.01.
